# Preferential Release of microRNAs via Extracellular Vesicles is Associated with Ductal Carcinoma In Situ to Invasive Breast Cancer Progression

**DOI:** 10.1101/2024.12.09.627283

**Authors:** Sugantha Priya Elayapillai, Samrita Dogra, James Lausen, Matthew Bruns, Fariba Behbod, Cole Hladik, Amy Gin, Chao Xu, Roy Zhang, Wei-Qun Ding, Bethany N. Hannafon

## Abstract

Ductal carcinoma in situ (DCIS) is a benign “pre-cancer” that increases the risk of invasive breast cancer (IBC). Not all DCIS progress to IBC, and the primary factors driving progression remain unclear. Small extracellular vesicles (sEVs) or exosomes are known to play a role in advanced cancers, but their involvement in DCIS is poorly understood. This study examined the role of sEVs and their RNA content in DCIS progression. Rab27A, which regulates exosome release, is elevated in DCIS and IBC tissues compared with normal breast tissues. Inhibition of sEV release via Rab27A knockdown alters pro-invasive pathways and reduces invasion in a DCIS mouse model. Using the isogenic MCF10 breast cancer progression series, we found a significant increase in microRNAs (miRNAs) in sEVs from normal to malignant states, with the highest number of differentially expressed miRNAs in IBC sEVs compared with DCIS sEVs. In vivo, DCIS invasive progression elevated circulating sEV miRNA levels, which decreased upon Rab27A knockdown. Re-expression of miR-205, preferentially loaded into IBC sEVs, reduced proliferation, invasion, and EMT marker expression in DCIS cells. Combined Rab27A knockdown and miR-205 expression repressed TGF-β signaling, activated p38, and induced cell cycle arrest and cell death. These findings illustrate that sEVs and their miRNAs promote DCIS progression, and the reintroduction of miR-205 in DCIS cells can inhibit invasive progression.

## Introduction

Every year, approximately 50,000 women are diagnosed with ductal carcinoma in situ (DCIS), a benign and nonobligate precursor of invasive breast cancer (IBC). [1] IBC is thought to arise from DCIS; however, not all DCIS become IBC. [2] Strategies to identify which DCIS have the potential to become malignant versus those that remain indolent are important clinical questions. [3] The progression of DCIS to IBC involves neoplastic proliferation and genetic changes arising from the luminal epithelial cell layer of the mammary duct, ultimately leading to breakdown of the mammary ductal structure and invasion into the surrounding stroma. [4, 5] However, whether these events are primarily due to DCIS cell-intrinsic events or extrinsic signaling remains unclear. Current research has shown that the intrinsic factors present in DCIS, such as genomic and transcriptomic changes, are similar to those in IBC. [6–11] This suggests that extrinsic factors, such as signaling within the tumor microenvironment, may play a predominant role in the progression of DCIS. However, our understanding of the extrinsic signaling events that occur in the DCIS microenvironment and their role in promoting invasive progression of DCIS is limited.

Although the molecular mechanisms of breast cancer progression have been well studied, extrinsic signaling involving small endosome-derived extracellular vesicles (sEVs), also known as exosomes, and their contents in the early stages of breast cancer have not been well defined. Recent studies have shown that sEVs mediate cell-to-cell communication through autocrine and paracrine mechanisms by transporting biologically active molecules, such as lipids, proteins, and RNAs, including microRNAs (miRNAs), to the surrounding cells and into the circulation. [12] sEVs may play an important role in breast cancer development and progression. For example, under hypoxic conditions, breast cancer cells release more EVs than normal epithelial cells [13], and EVs can transport oncogenic factors that can remodel the tumor microenvironment and contribute to breast cancer. [14–18] The small GTPase Rab27A regulates multivesicular endosome membrane docking at the plasma membrane and sEV/exosome secretion [19, 20] and may promote primary tumor growth. [21] We and others have previously demonstrated that Rab27A knockdown in breast cancer cells reduced sEV secretion. [21, 22] In these studies, depletion of Rab27A resulted in a reduction of miRNA sEV secretion [22], primary tumor growth, and lung metastasis due to modifications to the tumor microenvironment [21], indicating the importance of sEVs in cancer progression. miRNAs influence gene expression by regulating the expression of messenger RNAs (mRNAs), and are a substantial component of the RNA found in sEVs. Depending on the function of their target genes, miRNAs may regulate the expression of oncogenes and tumor suppressors. Many genetic events required for the invasive progression of DCIS occur at the pre-invasive stage, and these events include changes in miRNAs. [5] Aberrant expression can influence specific oncogenic tumor-suppressor pathways that are required for breast cancer progression. For example, miRNAs in DCIS cells can influence hormone signaling, cell-cell adhesion, epithelial-to-mesenchymal transition (EMT), TGF-β signaling, maintenance of cancer stem cells, and modulation of the extracellular matrix. [23] However, to our knowledge, the role of sEV signaling in the invasive progression of DCIS has not yet been described.

In this study, we sought to determine the role of sEVs in DCIS invasive progression and define the sEV RNA content at each successive stage of breast cancer development using the MCF10 isogenic breast cancer progression series as a model. We also characterized the role of sEVs and their specific miRNA content using in vitro and in vivo DCIS and IBC models. We demonstrated that circulating sEV miRNA levels change with disease progression and when invasive progression is attenuated, providing insights into the potential application of sEVs miRNAs as indicators of malignancy. We demonstrated that sEVs promote invasive programs and that specific miRNAs enriched in sEVs are preferentially released from DCIS and invasive breast cancer cells. We further demonstrated that inhibition of sEV release and re-expression of certain miRNAs can attenuate pro-invasive programs.

## Materials and Methods

### Cell lines and culture conditions

The untransformed normal cell line MCF10A and invasive carcinoma cell lines MCF7, ZR75-1, BT474, SKBR3, MDA-MB-231, and BT20 were purchased from the American Type Culture Collection (ATCC, Manassas, VA, USA) and cultured according to ATCC recommendations. The benign proliferative, MCF10AT1, and the invasive carcinoma stage MCF10CA1a were purchased from The Barbara Ann Karamanos Cancer Institute (Detroit, MI). The ductal carcinoma in situ (DCIS) cell lines SUM-102.gfp (SUM-102) and SUM-225.gfp (SUM-225) were a gift from Dr. Stephen P. Ethier (Medical University of South Carolina). Cells were cultured in accordance with the conditions provided by Dr. Ethier. [24] The DCIS cell line, MCF10DCIS.com (MCF10DCIS), was a gift from Dr. Fariba Behbod (University of Kansas Medical Center) and cultured in DMEM/F12, supplemented with 10 mM HEPES, 5% horse serum, 0.029 M NaHCO_3_, 100 IU/ml penicillin and 100 μg/ml streptomycin. The cells were maintained in a humidified chamber at 37°C and 5% CO_2_, except for SUM-102.gfp and SUM-225CWN.gfp cells, which require 10% CO_2_. The sEV-depleted serum was prepared by ultracentrifugation at 100,000 × g for 16 h. All cell lines were verified by STR analysis and were routinely checked for mycoplasma contamination.

### sEV isolation

sEVs were isolated from the culture medium using a combination of centrifugation, ultracentrifugation, and filtration as previously described. [22, 25] sEVs were isolated from mouse plasma samples using the Exoquick reagent (System Biosciences, EXOQ5A) following the manufacturer’s protocol. The resultant sEV pellet was suspended in 1X PBS and the sEV concentration was estimated using BCA Protein Assay (Pierce, 23225). Human CD63+ sEVs were enriched from mouse plasma samples using a magnetic bead-based immunoaffinity isolation method and CD63 antibody (Santa Cruz Biotechnology, sc-5275), as previously described. [25]

### Immunoblot analysis

Protein lysates were prepared by resuspending cells and sEVs in MPER buffer (78501, Thermo Fisher Scientific, Waltham, MA, USA) containing a Halt Protease and Phosphatase Inhibitor Cocktail (78442; Thermo Fisher Scientific). The sEV protein concentration was determined using the BCA Protein Assay (Pierce, 23225). Approximately 30–40 μg of protein from each sample was separated under reducing or non-reducing conditions for CD63 detection on a 12% SDS-PAGE gel and transferred to a polyvinylidene fluoride membrane. Specific protein levels were detected using primary and horseradish peroxidase-conjugated secondary antibodies. Immunoblots were analyzed using the Bio-Rad ChemiDoc Station.

### Nanoparticle tracking analysis

Isolated sEVs were suspended in PBS and stained with PKH67 Green Fluorescent Cell Linker (Sigma-Aldrich, MINI67, St. Louis, MO) by combining 1 µL of dye with 50 µL of diluent C; 10 µL of this solution was added to 20 µL of the sEV suspension and incubated for 5 min in the dark at RT. The stained sEV solution (25 µL) was diluted in 1.25 mL of distilled water and analyzed using a Nanosight NS300 System (Malvern Instruments, United Kingdom). Nanoparticles illuminated by the laser and their movement under Brownian motion were captured for 60 s. The videos were analyzed using Nanosight Tracking Analysis (NTA) software to determine the particle concentrations and size distribution profiles. Measurements were performed in triplicate for each sample. The size distribution and concentration profiles were averaged across replicates to derive representative size distribution profiles. Particle counts were normalized to the average cell count to derive the average number of particles released per 10^6^ cells.

### RNA sequencing

Total RNA from sEVs isolated from the conditioned medium of MCF10 cell line series or cultured cells was prepared using TRIzol reagent (Thermo-Fisher), followed by column clean-up using the Monarch RNA Cleanup kit according to the manufacturer’s protocol for whole transcriptome isolation (New England Biolabs, T2030S). RNA quality was checked using an Agilent 2100 Bioanalyzer, and concentrations from ThermoFisher’s NanoDrop readings were used.

#### Small RNA sequencing

Small RNA libraries were constructed using the New England Biolabs (NEB) NEBNext Multiplex Small RNA Library Prep Set for Illumina sequencing and the NEB standard protocol, as described previously. [25] Each library was indexed to four multiplex samples per sequencing run on the Illumina MiSeq platform using MiSeq 50 cycle Reagent Kits v2. The ExceRpt small RNA sequencing pipeline was used to analyze the sEV small RNA sequencing data. The sequencing (FASTQ) files were uploaded to Genboree.org and processed. Briefly, adapter sequences were auto-detected and trimmed, and endogenous reads were aligned using the following databases: miRNAs (miRbase, version 21), tRNA (gtRNAdb), piRNA (pIRNABank), Ensemble transcripts (Gencode, version 24), and circular RNA (circBase).[26] Raw aligned reads were normalized and analyzed using the R package DESeq2. [27] RNAs with zero counts in more than six out of 12 samples and total counts across all samples less than 10 were excluded owing to the high missing rate and limited information provided. Multiple testing was performed using Benjamini- and Hochberg-corrected P-values. The significance level was set at p < 0.05. T-distributed stochastic neighbor embedding (t-SNE) [28], and hierarchical clustering analyses were applied to present the sample similarity. Venn diagrams were constructed using InteractiVenn. [29]

#### Total RNA sequencing

Stranded RNA-seq libraries were constructed using the NEBNext Poly (A) mRNA isolation kit, followed directly by the IDT XGen Broad Range RNA Library Prep Kit and the established protocols. Library construction was performed using up to 1 µg of RNA. Each library was indexed during library construction for multiplex sequencing, and the quality and quantity were determined using Invitrogen’s Qubit 4 fluorometer and an Agilent 2100 Bioanalyzer. The samples were normalized and pooled onto a 150 paired end run on an Illumina NextSeq 2000 platform to obtain 20M reads per sample. The fastq files were aligned with the Human GRCh38 genome, and differentially expressed genes were identified using Illumina’s DRAGEN v4.2.4 app at Basespace. The output files included differential expression reports (genes.res), raw gene counts, heat maps, PCA plots, dispersion plots, and MA plots produced using DESeq 2 at the gene level for each comparison.

### Oncomine database, miRNA target prediction, and pathway analyses

The mRNA expression data for pre-invasive breast cancer were examined by querying the Oncomine cancer microarray and gene expression databases. [30] miR-205 levels in primary breast tissues were examined by querying the UALCAN database [31], and miRNA target mining and pathway analyses were conducted using the web-based tools miRWalk (accessed August 2021) [32] and Enrichr (accessed July 2024). [33]

### Quantitative RT-PCR (qRT-PCR)

*miRNA* - cDNA was synthesized from 100 ng of total RNA using a qScript miRNA cDNA Synthesis Kit (Quanta BioSciences) For miRNA qRT-PCR, an aliquot of cDNA, equivalent to 4 ng of the original RNA, was mixed with Perfecta SYBR Green SuperMix, Quanta Universal PCR Reverse Primer (Quanta BioSciences), and a sequence-specific forward primer or positive control forward primer (Quanta, Lot# 015644) in 20 μL PCR reactions. Synthetic *Caenorhabditis elegans* miR-54 (cel-miR-54) RNA oligonucleotides (Integrated DNA Technologies, Coralville, IA, USA) were spiked into RNA samples as control [34].

*mRNA* - 250ng of total RNA was converted to cDNA using an Iscript cDNA Synthesis Kit (Bio-Rad, 1708891). GAPDH was used as an internal control for mRNA expression analysis. The PCR products were amplified using a Bio-Rad CFX 96 Real-Time PCR (Bio-Rad, Hercules, CA, USA) instrument under the following conditions: 95°C for 2 min; 40 cycles of 95°C for 5 s, 60°C for 15 s, and 70°C for 15 s. Relative changes in gene expression were calculated using the ΔΔCT method. [35]

### Rab27A knockdown

The pLenti-CMV-Puro-shRNARab27A targeting vector was transduced with lentivirus into MCF10DCIS cells, as previously described. [22] Approximately 48 h post-infection, the cells were selected for gene transfer by treatment with 1μg/ml puromycin for 4 days.

### Cell proliferation assay

Cells were seeded in a 96-well plate at a density of 2,500 cells/well in quadruplicate. The cells were cultured at 37°C and 5% CO_2_, and cell growth for 1-4 days using an Incucyte live-cell imaging system. Live cell images were obtained using a 10X objective lens, and four images per well and percentage confluence were analyzed using Incucyte 2022 B software. At the end of the assay, cell proliferation was determined by colorimetric assay using 20 μL of CellTiter 96® AQueous One Solution (Promega, G3581) for 1h. The absorbance was recorded at 490 nm using a spectrometer.

### Invasion assay

MCF10DCIS cells were starved for at least 12 h before initiating the assay. Transwell inserts (8.0- μm pore size) were coated with 100 μg/ml Matrigel Basement Membrane Matrix (Corning, 354234). A 0.5 mL cell suspension of 2.5 × 10⁴ MCF10DCIS cells was added to the coated transwell/invasion chamber and incubated for 24-48 h in a humidified incubator at 37°C and 5% CO₂. The number of invaded cells was assessed by crystal violet staining or CyQuant cell proliferation assay. To detach cells, the invasion chamber was placed in a well containing 1X trypsin and incubated at 37°C for 15-20 min with shaking. Detached cells were assayed using a CyQUANT Cell Proliferation Assay Kit (Fisher Scientific, C7026) following the manufacturer’s protocol. The fluorescence signal was measured using a Bio-Tek Synergy H1 microplate reader with a 480/520 nm filter. Alternatively, cells in the Transwell insert were fixed with formalin, stained with 0.05% crystal violet in 20% methanol, and imaged using a Nikon Eclipse TE2000-U microscope at 10X magnification. After imaging, cells were destained with 10% glacial acetic acid, and absorbance was measured at 570 nm using a Bio-Tek Synergy H1 microplate reader. The number of invading cells was interpolated using linear regression analysis based on a standard curve of a cell suspension of known concentrations.

### Mouse-INtraDuctal transplantation Model (MIND)

Mouse experiments were conducted using an IACUC approved protocol at the University of Kansas School of Medicine. Cell implantation recipients were 8- to 10-week-old virgin female NOD-SCID IL2Rgamma^null^ (NSG) mice purchased from Jackson Laboratories. For MIND surgeries, a 30-gauge Hamilton syringe with a 50-μl capacity, and a blunt-ended 1/2-inch needle were used to deliver the cells, as previously described. [36, 37] Two microliters of PBS (with 0.04% trypan blue) containing 35,000 MCF10DCIS or MCF10DCIS+shRab27A cells were injected. For timed experiments, mice were euthanized at approximately 4-8 weeks (DCIS) and 8-10 weeks (invasive) post-transplantation as previously described. [36, 37] For shRab27A experiments, mice were euthanized at approximately eight weeks post-transplantation.

### Immunofluorescence staining

Tumor tissues were dissected from the mice upon sacrifice, fixed in 10% neutral buffered formalin solution, and embedded in paraffin. FFPE tissues were sectioned (4 µm) and mounted on positively charged slides. The slides were dried overnight at room temperature and incubated at 60°C for 45 min. Slides were transferred to a Leica Bond RX for dewaxing and then treated for target retrieval at 100°C for 20 min with a retrieval solution (pH 6.0) or pH 9.0. After permeabilization and blocking, the sections were incubated with primary antibodies for 60 min, followed by incubation with Alexa Fluor 568- or 647-conjugated antibodies for 60 min. The slides were quenched with TrueVIEW to reduce unwanted autofluorescence. DAPI was used as the counterstain. The slides were mounted using Vectashield Antifade Mounting Medium (Vector, SP-8400). Antibody-specific positive and negative controls (omission of primary antibody) were stained in parallel. Stained slides were scanned and analyzed using the Operetta High Content Imaging System (PerkinElmer) and associated Harmony Analysis Software.

### microRNA transfection

mirVana miRNA scramble negative control and miRNA mimics hsa-miR-30c-5p, hsa-miR-103a-3p, hsa-miR-205-5p, and hsa-miR-222-3p were purchased from Thermo Fisher Scientific. Lipofectamine^TM^ RNAiMAX transfection reagent (Thermo Fisher,13778150) in OptiMEM (Gibco) was used to transfect scramble or miRNA mimics, according to the manufacturer’s instructions.

### Cell cycle and apoptosis assay

MCF10DCIS WT and shRAB27A cells were transfected with scrambled or miR-205 mimics using the Lipofectamine RNAiMAX transfection reagent (Thermofisher,13778150). Cell cycle analysis was performed using the Click-iT Edu AF 488 flow cytometry assay kit (Thermo Fisher, C10632) and staining with PO-PRO-1 Iodide according to the manufacturer’s instructions. EDU incorporation was determined by flow cytometry. Apoptosis was analyzed using an annexin V apoptosis kit (Biotium, 30061), according to the manufacturer’s instructions. Annexin V and PO-PRO-1 Iodide were detected at 488 and 435 nm, respectively, using flow cytometry. A Stratedigm S1200Ex Flow cytometer was used, and the data were analyzed using ModFit Software

### TGF β/SMAD luciferase reporter assay

TGFβ/SMAD signaling activity was measured using the SBE4-luc reporter construct (Addgene, 16495) in MCF10DCIS WT and shRAB27 cells (1 × 10^5^ cells/well) in 12 well plate. Briefly, cells were co-transfected with 2 μg SBE4-luc reporter and 0.2 μg renilla luciferase vector (PRL-TK, Promega) using the X-tremeGENE 360 transfection reagent (Sigma, XTG360-RO). After 72 h of transfection, cell lysates were assayed according to the manufacturer’s protocol using the Dual-Luciferase Reporter Assay system (Promega, E1910). Luminescence was read using a Synergy H1 Microplate Reader (BioTek). TGFβ firefly luminescence values were normalized to Renilla luciferase luminescence values.

### Statistical analysis

Statistical analyses were performed using GraphPad Prism software (GraphPad Software). When appropriate, Student’s t-test for experiments and ANOVA with post hoc analysis for multiple comparisons were applied to determine significant differences between the control and experimental groups, with p <0.05, as the level of significance.

## Results

### Rab27A knockdown altered EMT in vitro and attenuated invasion in a mouse intraductal (MIND) xenograft model

To evaluate the contribution of sEVs to DCIS invasive progression, we first examined RAB27A expression in a gene expression dataset from the micro-dissected epithelia of normal, DCIS, and IBC tissues. [38] RAB27A expression increased 2.3-fold in DCIS (p=0.002) and 2.2-fold in invasive (p=0.0004) cells compared to that in normal cells (Figure 1A) [38], suggesting that RAB27A is elevated at the DCIS stage of breast cancer. To examine the role of sEVs in DCIS invasion, Rab27A was knocked down in MCF10DCIS (shRab27A) cells, as confirmed by immunoblotting (Figure 1B). sEV secretion was significantly reduced by 64-73% in shRAB27A knockdown cells relative to parental MCF10DCIS cells (Figure 1C). Although sEV secretion was reduced, cell proliferation was not significantly different between MCF10DCIS and shRab27A cells (Figure 1D). RNA sequencing analysis identified 697 differentially expressed genes between shRab27A and MCF10DCIS cells (fold-change ≥1.5, ≤1.5, adjusted p<0.05) (Figure 1E). Pathway analysis revealed enrichment of genes involved in EMT, extracellular matrix (ECM) receptor interaction, and apical junctions. (Figure 1F). As previous studies have shown that sEVs are major inducers of EMT [39–41], which plays a role in invasion, we validated the effect of Rab27A knockdown on EMT regulators. While no increase in E-cadherin (epithelial-state marker) was observed, vimentin (mesenchymal-state marker) expression was significantly decreased (p<0.05, p<0.0001, respectively). Similarly, SNAIL, ZEB1, ZEB2, and TGFB1 (p<0.05, p<0.01) decreased, most likely due to the significant decrease in the transcription factors FOXC2 and SNAIL (p<0.01 and p<0.05, respectively) (Figure 1G). Next, we used a Mouse INtraDuctal (MIND) transplantation model [36, 42] to evaluate the role of Rab27A and sEV secretion in invasion progression in vivo. MCF10DCIS or shRab27A cells were implanted into the mouse mammary ducts (Figure 1H). DCIS and invasive lesions were confirmed by histological staining of mouse mammary glands for the epithelial marker cytokeratin 5 (CK5) and the myoepithelial cell marker smooth muscle actin (SMA) (Figure 1I). The same number of DCIS lesions was observed in MCF10DCIS and shRab27A implanted mice, whereas the number of invasive lesions was significantly decreased in the shRab27A group (p<0.001) (Figure 1J), indicating that Rab27A is associated with invasive progression. Taken together, these data demonstrate that RAB27A-mediated sEV release alters programs associated with EMT, ECM, cell-cell adhesion in vitro, and invasive progression in vivo.

**Figure 1.**
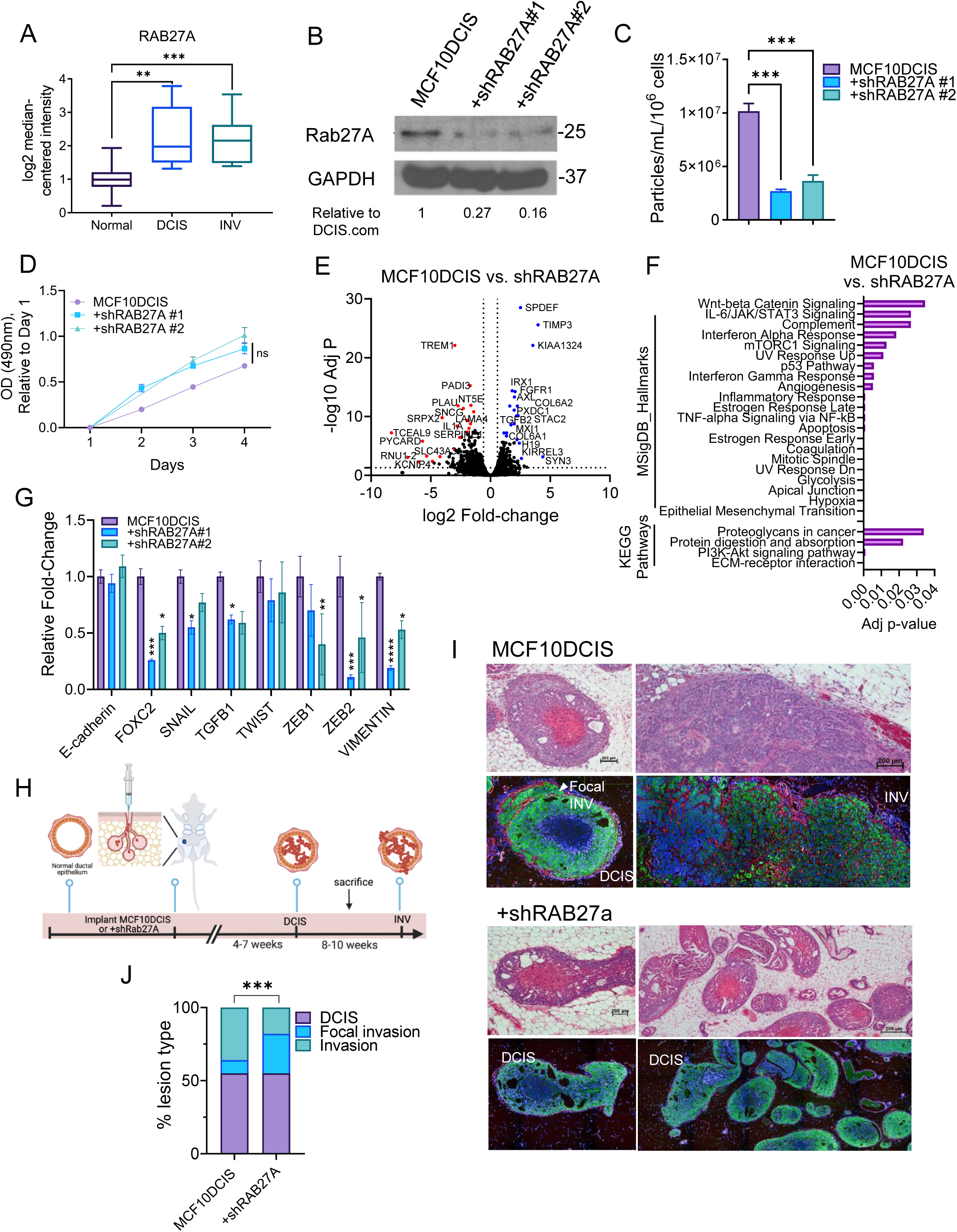
RAB27A knockdown alters expression of EMT factors in vitro and decreases DCIS to invasive progression in vivo. **A.** Boxplots of RAB27A expression in microdissected epithelia of normal mammary tissue, DCIS, and invasive breast cancer (INV) from Ma et al. study on Oncomine database. **B.** Immunoblot of RAB27A in MCF10DCIS and shRAB27A cells. GAPDH was used as a loading control. **C**. NTA of exosomes isolated from conditioned cell culture medium. Data are depicted as nanoparticles per mL per 10^6^ cells relative to MCF10DCIS cells (n=3). The results are presented as mean ± SEM (***p < 0.001, t-test). **D**. Growth curves of MCF10DCIS and +shRAB27A cells determined by MTS assay (n=3) relative to day 1. The results are presented as the mean ± SEM. **E**. Volcano plot of differentially expressed (increased, blue and decreased, red) genes in MCF10DCIS vs +shRab27A cells. The horizontal line indicates a significant p-value ≥1.3 log10 P value), and vertical lines indicate ≥1.5 fold-change (log2 fold change ≤ -0.58 or ≥0.58). **F.** Significantly enriched KEGG and Molecular Signature Database (MSigDB) Cancer Hallmarks pathways in MCF10DCIS vs shRab27A cells from E. **G**. qRT-PCR analysis of EMT-related transcription factors and downstream targets in MCF10DCIS and shRab27A cells. Data are shown relative to MCF10DCIS cells and normalized to 36B4 (n=3). Bars represent the mean ± SEM, \**p*<0.05, \*\**p*<0.01, ***p<0.001, ****p<0.0001, t-test. **H**. Schematic of mouse intraductal (MIND) model procedures **I**. Histological staining of DCIS (in situ) and invasive lesions in dissected mouse mammary glands. *Upper-*Hematoxylin and eosin (H&E) staining. The scale bar represents 200μm. Immunofluorescence (IF) staining of the myoepithelial cell marker, smooth muscle actin (SMA; red); luminal epithelial cell marker, cytokeratin 5 (CK5; green); and nuclei, DAPI staining (blue) in mice implanted with MCF10DCIS (upper) and +shRAB27A cells (lower). **J**. The percentages of DCIS, focal invasion, and invasive lesions were determined in the mouse mammary glands (n=6 mammary glands, 3 mice per group). ***p<0.001, Fisher’s Exact Test.

### sEVs are progressively enriched in miRNAs that are associated with breast cancer progression

To understand how the content of sEVs changes from benign to IBC and contributes to invasive programs, we isolated sEVs from each cell line in the MCF10 isogenic series comprising MCF10A (normal mammary epithelial cells), MCF10AT1 (benign proliferative cells), MCF10DCIS (spontaneous malignant progression), and MCF10CA1 (malignant cells with metastatic capabilities) [43], and profiled their small RNA content. sEVs were isolated and probed for classic exosome markers (Flotillin-1, Tsg101, CD63) and the cell-specific marker calnexin to confirm the isolation of sEVs. The size of the sEVs ranged from 100 to 250 nm (mean = 160 nm) and did not differ significantly among the MCF10 series, while the number of secreted sEVs increased with each progressive stage, except for MCF10AT1, which secreted fewer sEVs than the other lines. Next, we profiled the small RNA content of the sEVs. T-distributed stochastic neighbor embedding (t-SNE) analysis revealed four distinct sEV RNA clusters (Figure 2A). To evaluate the types of small RNAs present, reads were aligned to multiple small RNA databases. MCF10A sEVs were highly abundant in small nucleolar RNAs (snRNAs), constituting 75.37% of the small RNA content, whereas only 13.4% of the small RNA content comprised microRNAs (miRNAs). In contrast, miRNAs accounted for 76.67% of MCF10AT1, whereas snRNAs comprised only 5% of the sEV content of MCF10AT1, a 70% decrease from MCF10A. Increased miRNA content was also observed in MCF10DCIS and MCF10CA1 cells, at 83% and 86%, respectively (Figure 2B). On the basis of these results, we focused on the remaining studies on the miRNA content of sEVs.

**Figure 2.**
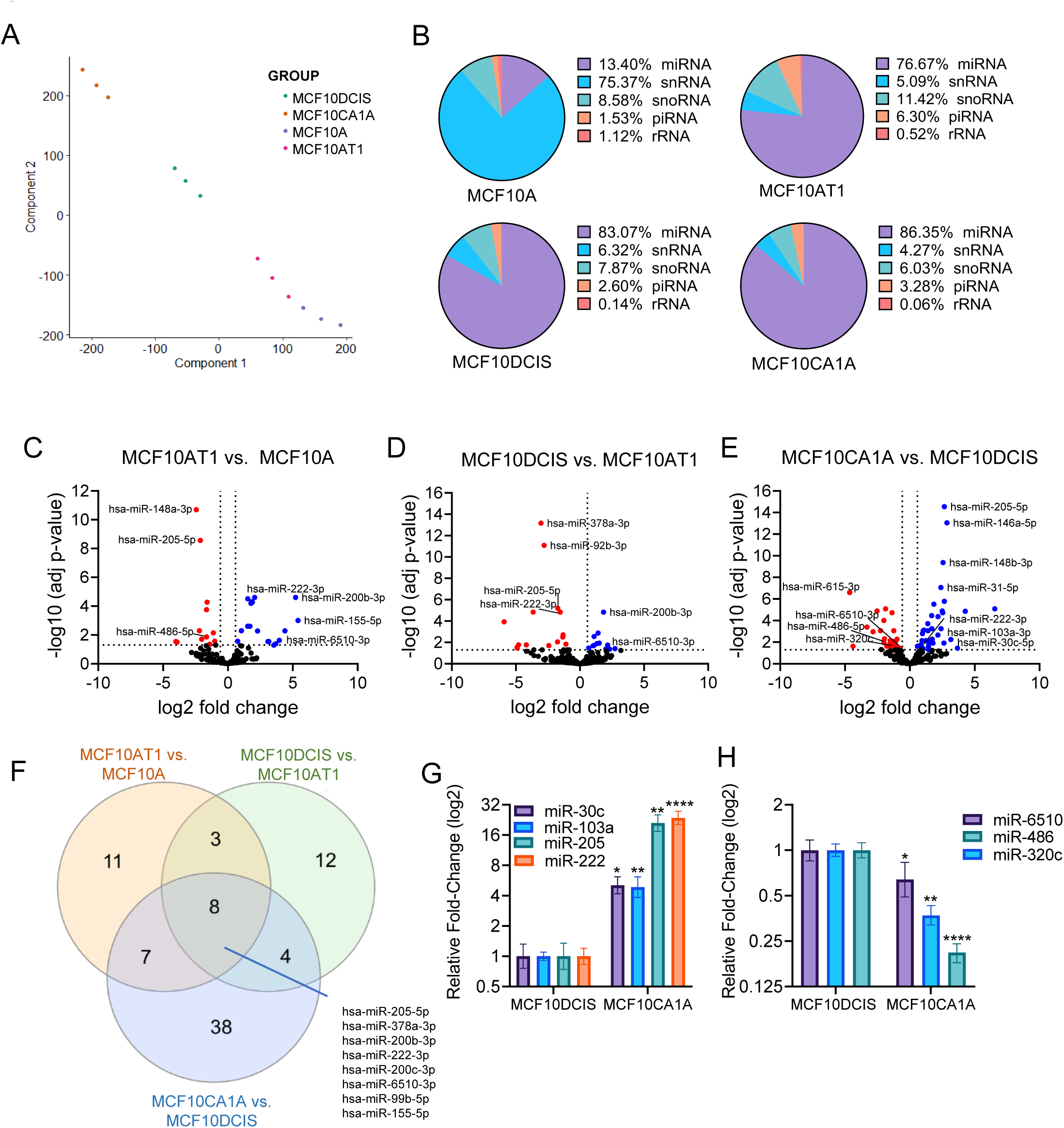
sEV RNA content is enriched with miRNAs associated with breast cancer progression. **A**. Sample similarity based on the aligned reads was determined by t-SNE analysis using the raw read counts of 95 significantly differentially expressed miRNAs among all groups. **B.** The proportional distribution of reads per million was detected among the following ncRNA types: ribosomal RNA (rRNA), piwi-RNA (piRNA), small nucleolar RNA (snoRNA), small nuclear RNA (snRNA), and microRNA (miRNA) in the MCF10 series. **C-E**. Volcano plots of quantitative differences in miRNA levels for each comparison. Red dots above the dashed line represent miRNAs with significant differences (false discovery rate [FDR] < 0.05). **F**. Venn diagram of sEV miRNAs significantly differentially expressed in each pair-wise comparison. qRT-PCR of sEV miRNAs normalized fold-change in expression levels relative to MCF10DCIS with increased expression **(G**) and decreased expression (**H**). Data are presented as mean ± SEM (n = 3). *p<0.05, **p<0.001, student t-test.

Hierarchical clustering analysis was performed on the normalized miRNA reads, revealing four moderately well-separated clusters of miRNAs (fold change ≥1.5, ≤1.5, and p≤0.05). Between MCF10AT1 and MCF10A, 29 miRNAs differed (12 down, 17 up), whereas 27 miRNAs differed between MCF10DCIS and MCF10AT (15 down, 2 up). The largest differential expression of miRNAs was observed in MCF10CA1A compared to MCF10DCIS, with 57 miRNAs (25 down, 32 up) (Figure 1C-E). Eight miRNAs were commonly altered in all comparisons, and a set of 38 miRNAs were distinct from the MCF10CA1 and MCF10DCIS comparisons, suggesting that these miRNAs could potentially be involved in the DCIS to invasive transition. miRNAs with high abundance (mean normalized counts >90) that were increased (miR-30c, miR-103a, miR-205, and miR-222) and decreased (miR-6510, miR-486, and miR-320c) in MCF10CA1 compared with MCF10DCIS were validated by qRT-PCR (Figure 1G-H). These data illustrate that sEVs are selectively enriched with miRNAs in the progression from benign to malignant states, and that the greatest change in miRNA expression occurs during the transition from DCIS to IBC.

### sEV miRNAs can be detected in the circulation and are altered by invasive progression in vivo

We previously reported that sEV miRNAs indicative of breast cancer can be detected in the plasma samples of human breast cancer patients and patient-derived orthotopic xenograft models. We then investigated whether sEV miRNAs could be detected in the plasma of mice implanted with MCF10DCIS or shRab27A cells (Figure 1H-1I). Plasma sEVs were isolated and enriched for human CD63 as previously reported. [25] Indeed, miR-30, -103a, -205, and -222 were detected in the plasma of MCF10DCIS implanted mice, and their levels were reduced compared to shRab27A implanted mice (Figure 3A). Based on these results, we speculated that changes in plasma sEV miRNAs may occur during invasive progression. Using a timed progression MIND model, we collected mammary glands and plasma from mice to 4-8 weeks (DCIS) and 8-10 weeks (invasive) post-injection of MCF10DCIS cells (Figure 3B). DCIS and invasive lesions were histologically confirmed (Figure 3C). sEVs were isolated from mouse plasma and were enriched for human CD63. The sEV protein concentration, as a proxy for the number of sEVs, was significantly higher (p<0.01) in mice bearing invasive tumors than in mice bearing DCIS (Figure 3D). miR-103a and miR-222 were significantly increased (p<0.05) and miR-486 was significantly decreased (p<0.05) with disease progression (Figure 3E), consistent with the RNA sequencing results. These results illustrate that miRNA levels in circulating sEVs are reflective of disease state.

**Figure 3.**
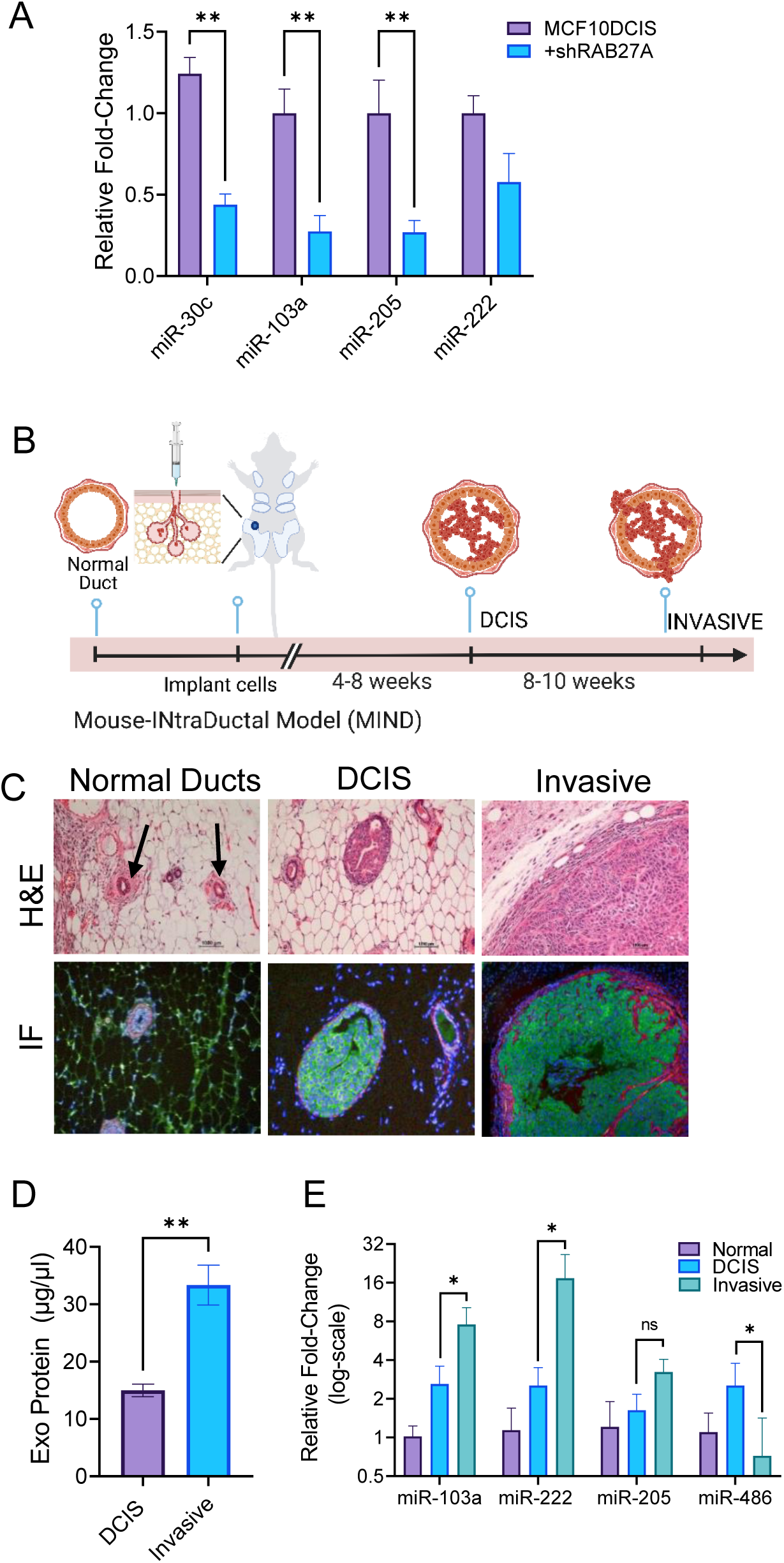
Circulating sEV miRNAs are altered in DCIS to invasive progression xenograft model. **A.** qRT-PCR analysis of sEV miRNAs in MCF10DCIS and shRab27A cells. Normalized fold-change in expression levels in invasive cells is shown relative to MCF10DCIS. The results represent mean ± SEM, n=3, *p<0.05, Student’s t-test. **B.** Schematic of the procedures involved in the intraductal injection of MCF10DCIS and an approximate timeline of DCIS and invasive development; created with BioRender.com. **C**. Normal mammary glands, DCIS, and invasive lesions were confirmed by histological staining of dissected mammary glands. Upper-Hematoxylin and eosin (H&E) staining. The scale bar represents 1000 μm. Immunofluorescence (IF) staining of the myoepithelial cell marker, smooth muscle actin (SMA; red); luminal epithelial cell marker, cytokeratin 5 (CK5; green); nuclei, DAPI staining (blue). **D.** Quantitation of circulating sEV protein concentration in plasma samples. Data represent mean ± SEM; **p<0.001, Student’s t-test. **E.** qRT-PCR analysis of enriched human CD63+ plasma sEV miRNA expression. Normalized fold-changes in expression levels in invasive tissues are shown relative to those in DCIS and normal tissues. The results represent mean ± SEM, n=3, *p<0.05, Student’s t-test.

### sEV-enriched miRNAs reduce proliferation and invasion and alter EMT transcriptional programs

Because miRNAs can reduce mRNA stability and translation through sequence-specific targeting, we performed miRNA target prediction and pathway enrichment analysis to identify genes and pathways potentially affected by miRNAs enriched in sEVs. Indeed, the miRNA targets in all comparisons were enriched for “pathways in cancer” and “signaling pathways regulating pluripotency of stem cells”.

Next, we focused on the miRNAs miR-205, miR-222, miR-103, and miR-30c, which were significantly increased in MCF10CA1A cells compared to MCF10DCIS cells, validated by qRT-PCR, and are established regulators of EMT (Figure 1G). [44–47] miRNAs can be preferentially retained in cells or released into sEVs. [48] To understand the cellular retention or release characteristics of these miRNAs, their levels were measured in MCF10DCIS and shRab27A cells, and sEVs. miR-103 and miR-30c levels were significantly lower (p<0.05) in sEVs than in cells, suggesting preferential cellular retention. In contrast, miR-205 and miR-222 levels were significantly higher in sEVs than in cells (p<0.05 and p<0.0001, respectively), suggesting preferential release. miR-30c, -103a, and -205 levels were significantly reduced in shRab27A sEVs relative to MCF10DCIS sEVs (p<0.05, p<0.001), suggesting that Rab27A knockdown resulted in less miRNA loading in the sEVs (Figure 4A). When we compared the cellular levels of miR-205 and miR-222, they were significantly increased (p<0.01) in shRAb27A cells compared to MCF10DCIS cells, demonstrating that knockdown of Rab27A inhibited their release, resulting in their cellular accumulation (Figure 4B).

**Figure 4.**
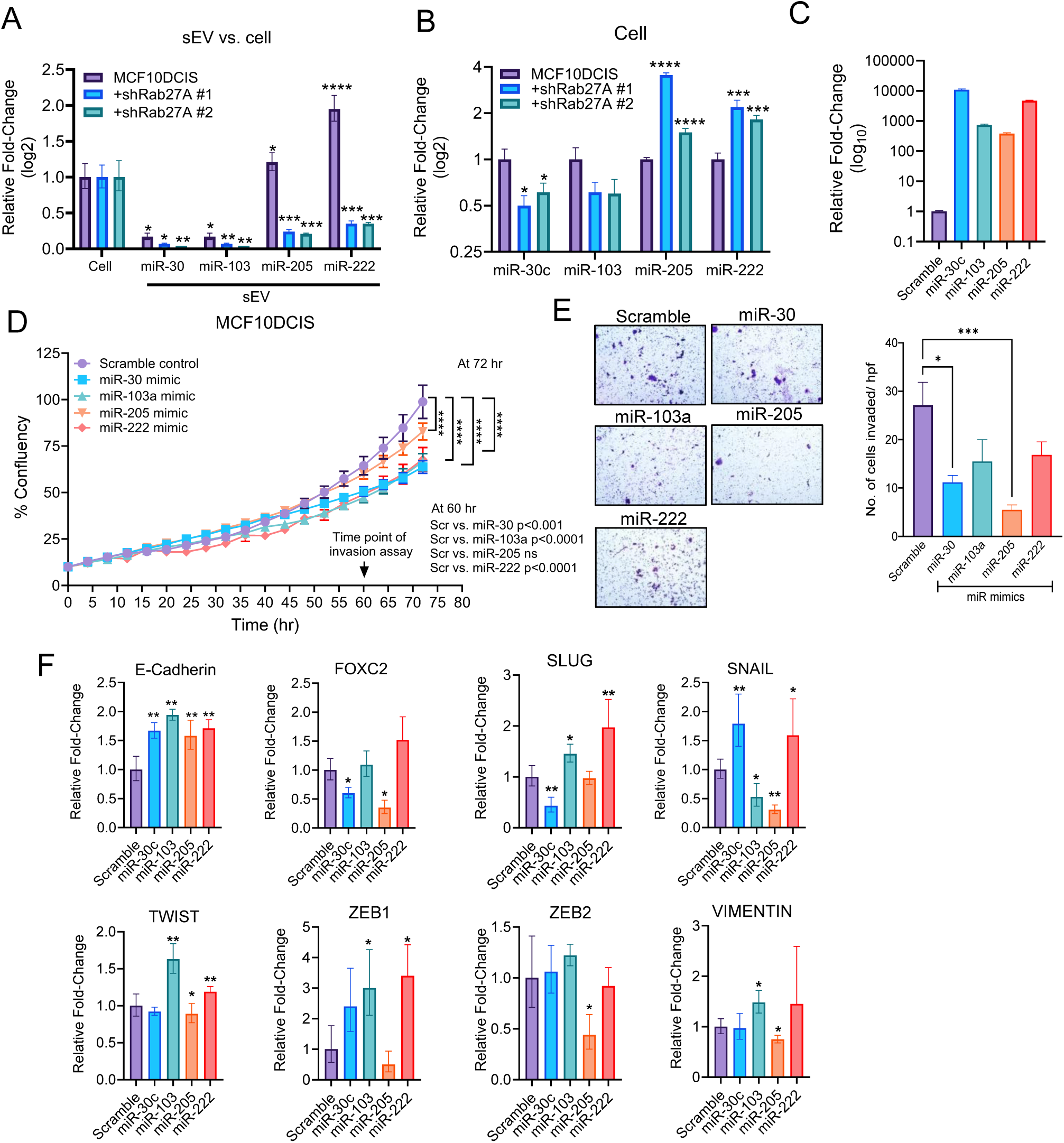
sEV-enriched miRNAs reduce proliferation and invasion and alter EMT transcriptional programs. qRT-PCR analysis of sEV-enriched miRNAs in sEVs relative to cellular levels (**A**) and MCF10A cellular levels compared with shRab27A cellular levels (**B**). Results represent the mean ± SEM, n=3, *p<0.05, **p<0.01, **p<0.001, ***p<0.0001, student t-test. **C**. qRT-PCR of miRNA mimic expression levels in MCF10DCIs cells. **D**. Proliferation of MCF10DCIS cells transfected with miRNA mimics or scrambled control was measured using the Incucyte system (n=12). The results are presented as mean ± SEM. ****p<0.0001, two-way ANOVA with post hoc test. **E**. Invasion of MCF10DCIS cell transfected with miRNA mimics or scramble control through the matrix in trans-well culture. The cells were allowed to migrate across the inserts for 48 h. The results represent the mean ± SEM, *p<0.05, ***p<0.001, ANOVA. **F**. qRT-PCR of EMT factors in MCF10DCIS cells transfected with miRNA mimic or scramble control.

Next, we sought to determine the individual contributions of each miRNA to the regulation of cell proliferation, invasion, and EMT. Using miRNA mimics, we overexpressed miR-205, miR-222, miR-103, and miR-30c in MCF10DCIS cells and confirmed their increased levels relative to those in the scrambled control (Figure 4C). Cellular proliferation was significantly reduced (p<0.0001) in all cells transduced with the miRNA mimics at 72 h (Figure 4D). Invasion was significantly reduced in miR-205 (p<001) and miR-103a (p<0.05) transduced cells relative to the scramble control (Figure 4E). The expression of E-Cadherin was significantly increased by all the miRNAs (p<0.01), while miR-30c reduced two of the mesenchymal factors (FOXC2 and SLUG) (p<0.05), and miR-205 consistently resulted in a significant decrease in expression of FOXC2 p<0.05), SNAIL (p<0.01), TWIST (p<0.05), ZEB2 (p<0.05), and VIMENTIN (p<0.05) relative to the scramble control (Figure 4F).

### miR-205 is elevated in breast tumors and preferentially loaded into sEVs in invasive breast cancer cells

Given miR-205’s ability to decrease cell proliferation and invasion, influence EMT transcriptional programs, and its preferential release in sEVs from MCF10DCIS cells, we aimed to explore its role further. Using the MCF10A progression model, we observed that miR-205 levels in sEVs increased relative to cellular levels at each stage, with the highest concentrations found in sEVs from invasive cells (MCF10CA1A) (p<0.05) (Figure 5A). We extended this analysis to include two additional DCIS cell lines, SUM-102 and SUM-225, and six IBC cell lines representing various breast cancer subtypes. Compared to MCF10A cells, cellular miR-205 expression was reduced in both SUM cell lines and further diminished in all invasive lines, whereas sEV miR-205 expression was decreased in both SUM lines but consistently elevated in sEVs from all invasive cell lines (Figure 5B and 5C). Examination of the Cancer Genome Atlas Data (TCGA) confirmed that miR-205 expression was significantly and consistently reduced in primary breast tumors compared to normal breast tissues, regardless of the pathological subtype (Figure 5D). These data demonstrate that miR-205 expression is lost in invasive breast cancer tissues, likely due to the preferential loading of miR-205 into sEVs from IBC cells.

**Figure 5.**
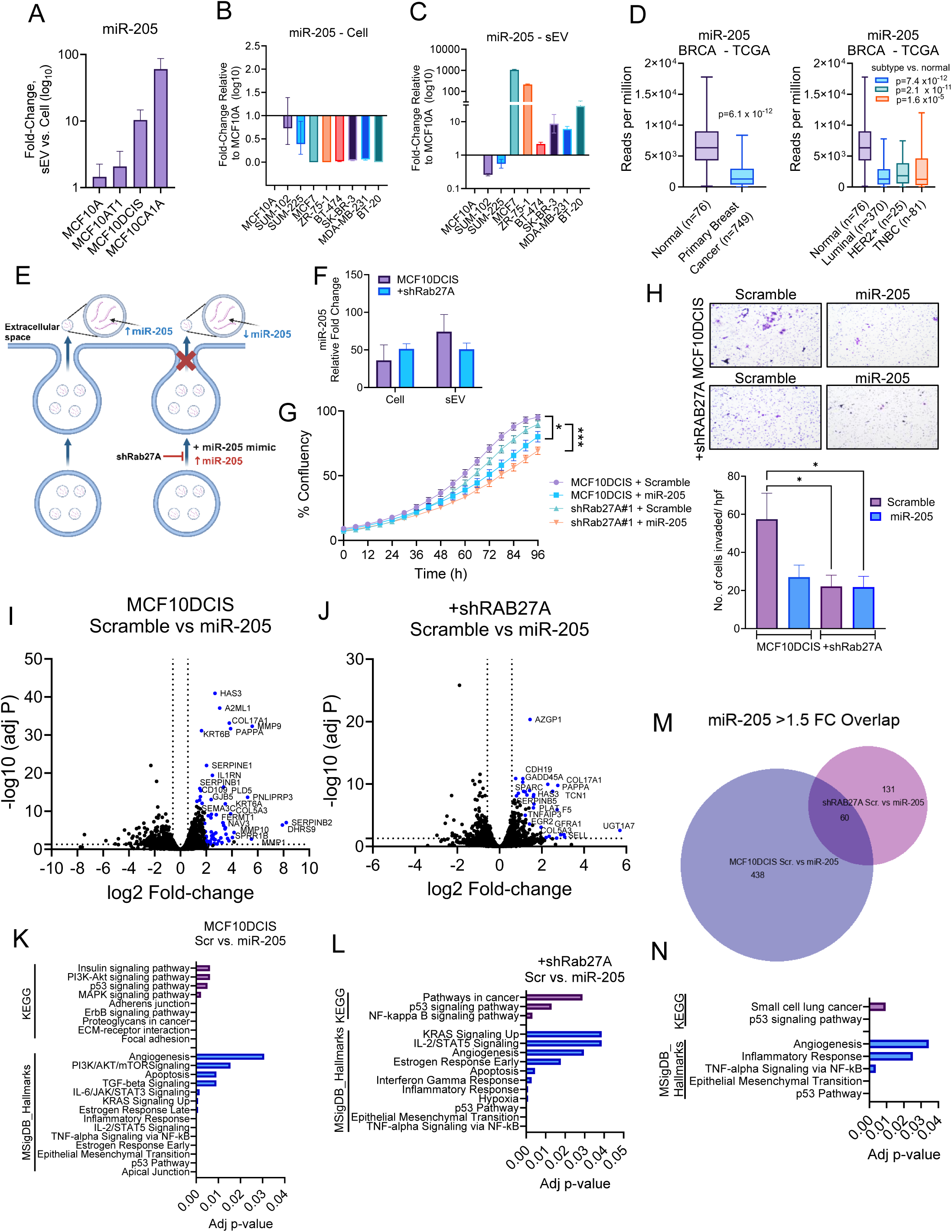
miR-205 is enriched in sEVs from invasive breast cancer cells and miR-205 over-expression combined with Rab27A knockdown reduces proliferation, invasion, and altered pathways related to invasion and stress response. **A**. qRT-PCR analysis of miR-205 in MCF10A progression series in sEVs compared to cellular levels. The results represent the mean ±_SEM, n=3, *p<0.05, Student’s t-test. qRT-PCR of miR-205 in DCIS (SUM-102, SUM-225), invasive breast cancer cells (**B**), and sEVs (**C**) relative to levels in MCF10A cells. The results are presented as mean ± SEM, n=3. **D**. Box-Whisker plot of miR-205 expression in breast cancer (BRCA), data from the Cancer Genome Atlas (TCGA), and P-values determined using student t-test. **E**. Schematic of the effect of miR-205 overexpression combined with Rab27A knockdown. **F.** qRT-PCR analysis of miR-205 expression. **G.** Cell proliferation as a measure of % confluency over time. **H**. Transwell invasion assay. Representative crystal violet-stained transwells and quantitative measurement of the number of cells invaded relative to the scramble control. Data represent mean ± SEM, n=9, *p<0.05, Student’s t-test. Volcano plot of significant (adj p<0.05) differential expression (>1.5 fold-change) in MCF10DCIS (**I**) and MCF10DCIS +shRab27A cells transfected with scramble miRNA or miR-205 mimic. Significantly (adj p<0.05) enriched KEGG and MSigDB pathways in scrambled compared with miR-205 transfected MCF10DCIS (**K**) and +shRab27A cells **(L**). Venn diagram of significantly differentially expressed genes (M) and significantly enriched pathways of overlapping genes (**N**).

### miR-205 overexpression and Rab27A knockdown results in reduced proliferation, invasion, and altered pathways related to MAPK, EMT, ECM-receptor, and p53 signaling

We then investigated the impact of sEV release of miR-205 on cellular programs by inhibiting its release through Rab27A suppression and constitutive miR-205 overexpression (Figure 5E), as confirmed by qRT-PCR (Figure 5F). The effects of miR-205 overexpression on the proliferation and invasion of MCF10DCIS and shRab27A cells were evaluated. miR-205 significantly reduced cell proliferation in MCF10DCIS (p<0.05), which was further decreased in shRAb27A cells (p<0.001) (Figure 5G) compared to scramble control. Both miR-205 overexpression and Rab27A knockdown, alone and in combination, significantly reduced invasion, although addition of miR-205 did not further decrease invasive activity in shRab27A cells (Figure 5H).

RNA sequencing was conducted to evaluate the gene expression programs modulated by increased miR-205 expression and Rab27A knockdown. Approximately 938 genes showed significant differences between the scrambled control and miR-205 overexpression in MCF10DCIS cells, with 498 upregulated in scrambled cells compared to miR-205 (Figure 5I). Fewer genes differed significantly (approximately 424) between +shRab27A cells with scramble and miR-205, with 191 upregulated in scramble compared with miR-205 (Figure 5J). Pathway analysis of the differentially expressed genes due to miR-205 expression in MCF10DCIS and shRab27A cells showed enrichment for MAPK and TGF-β signaling, ECM-receptor interaction, focal adhesion, EMT, and p53 pathway signaling (Figure 5K and 5L). A comparison of the genes modulated by miR-205 in MCF10DCIS and shRab27A cells revealed 60 overlapping genes between the datasets (Figure 5M). Pathway analysis of these 60 genes demonstrated enrichment of the p53 signaling pathway and EMT (Figure 5N).

### miR-205 regulates TGFβ and p38 MAPK signaling to induce G1 cell cycle arrest and apoptosis

TGFβ has a biphasic role in cancer, functioning as a tumor suppressor in the early stage of cancer and a promoter at later stages. [49] We observed that miR-205 overexpression reduced proliferation, invasion, and EMT factor expression (Figure 5) and that the TGF-β signaling pathway was altered by miR-205 overexpression (Figure 5K). Therefore, we first evaluated the role of miR-205 in TGF-β signaling using the TGFβ/SMAD reporter assay. We found that miR-205 overexpression significantly reduced TGF-β/SMAD signaling in both MCF10DCIS and +shRab27A cells (Figure 5A). We also observed altered p38 MAPK and p53 signaling with miR-205 overexpression (Figure 5K–5N), and downregulation of TGFβ signaling was associated with the activation of stress-activated protein kinases, such as p38, leading to cytotoxic cell death. [50] Therefore, we examined the effect of miR-205 overexpression, in combination with Rab27A knockdown, on this pathway. A >2-fold increase in phospho-p38 (Thr180/Tyr182) was observed in both MCF10DCIS and +shRAb27A cells overexpressing miR-205 (Figure 5B). G1 cell cycle arrest was associated with increased phospho-p53 (Ser15) levels in +shRab27A cells transfected with miR-205 (Figure 6E) and increased p21 levels in both MCF10DCIS and +shRAb27A cells transfected with miR-205 (Figure 6F), suggesting activation of the p53 response and p21, leading to the observed G1-phase arrest (Figure 6C, 6D). Next, we examined apoptotic and necrotic cell death and found that reduced Rab27A resulted in increased early apoptosis, whereas the addition of miR-205 increased the number of necrotic cells (Figure 6G). These results suggest that the combination of shRab27A and miR-205 alters the pathways that result in G1 phase cell cycle arrest and necrotic cell death through the activation of p38 MAPK, p53, and p21.

**Figure 6.**
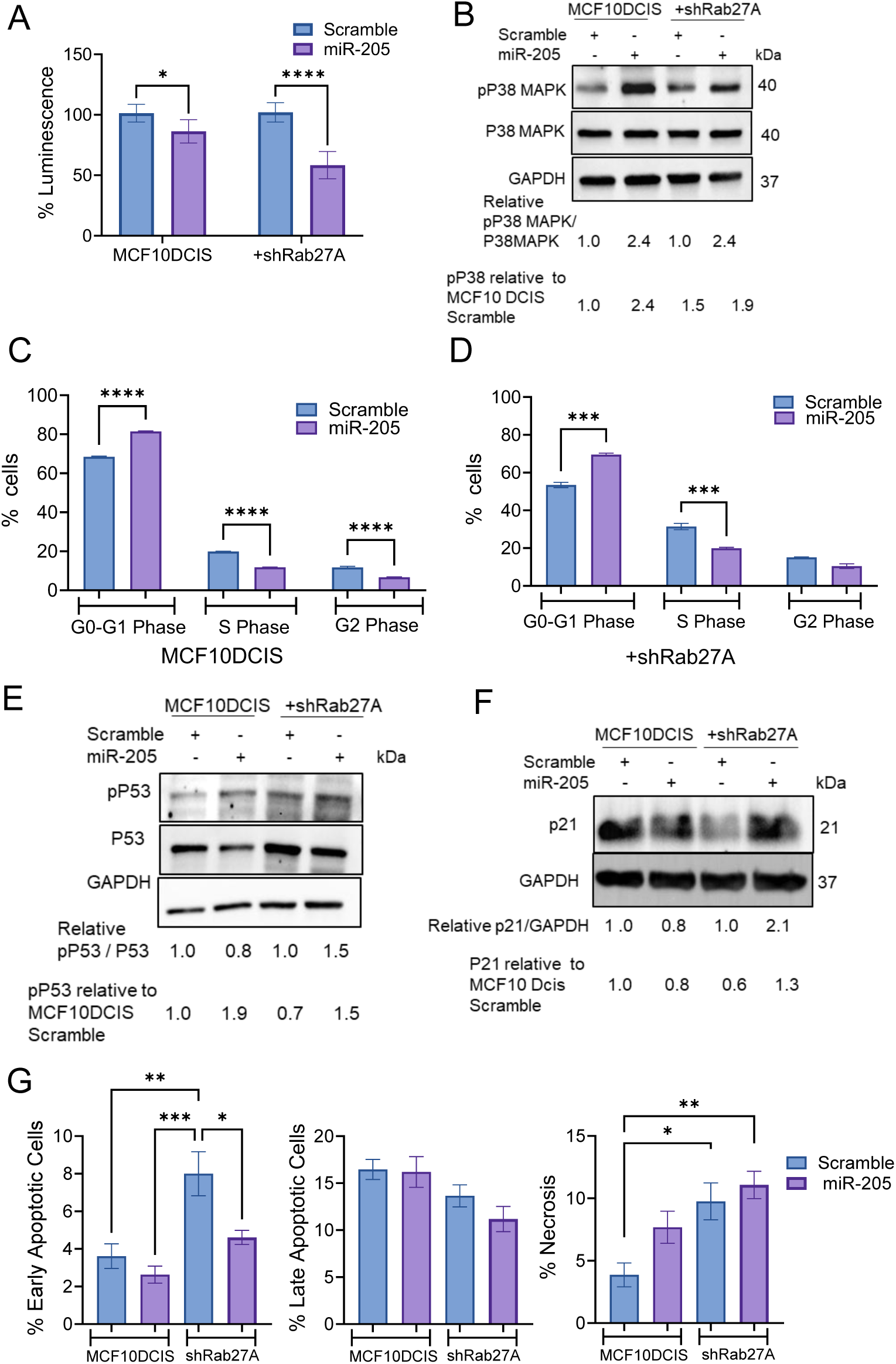
Combined Rab27A knockdown and miR-205 re-expression inhibited TGF-β signaling and activated p38, inducing cell cycle arrest and apoptotic death. **A.** TGF-β/SMAD reporter assay, data represent mean ± SEM *p<0.05, ***p<0.001, ***p<0.001, student t-test. **B**. Immunoblots of pP38 and P38. GAPDH was used as a loading control. Cell cycle phase assessment in MCF10DCIS (**C**) and shRab27A cells (**D**) were treated with scrambled miRNA or miR-205 mimic. Data represent the mean ± SEM; ***p<0.001, ****p<0.0001, Student’s t-test. Immunoblotting for pP53, P53 (**E**) and p21 (**F**). GAPDH was used as a loading control. **G**. Flow cytometric analysis of apoptosis and necrosis. The percentages of cells with early and late apoptosis and necrosis are shown. Data are presented as mean ± SEM. *p<0.05, **p<0.01, ***p<0.001, one-way ANOVA.

## Discussion

While numerous studies investigating the role of sEVs in cancer have been reported, the majority of breast cancer sEV-related research has focused on later events in cancer progression, such as their effect on primary tumor growth and metastasis or paracrine signaling between different cell types within the tumor microenvironment. [51] To the best of our knowledge, this is the first study to examine the role of sEVs and their RNA content in early breast cancer progression and to evaluate the role of sEVs in promoting invasive progression in DCIS models both in vitro and in vivo.

In this study, the MCF10A isogenic cell line series was used to characterize the changes in sEV content within each defined progressive stage of breast cancer. The primary observation was that in H-Ras-transformed cells (MCF10AT1), the small RNA content of their sEVs was significantly altered compared to that in spontaneously immortalized but non-transformed MCF10A cells. The miRNA content of sEVs from H-Ras-expressing MCF10AT1 cells increased by 64% compared to that of MCF10A sEVs. Previous studies have examined the role of Ras oncogenes in sEV signaling and content loading. For instance, sEVs from H-Ras-transformed Madin-Darby Canine Kidney cells were programmed to contain cancer-promoting proteins, such as matrix metalloproteinases, integrins, and RNA-binding proteins. [52] In another investigation, H-Ras and K-Ras signaling in prostate cancer cells resulted in sEVs with abundant oncogenic miRNAs. [53] In isogenic colon cancer cell lines expressing either wild-type or mutant K-Ras, differences in sEV miRNA profiles were observed in the K-Ras mutants relative to the parental cells, indicating a K-Ras-dependent sorting mechanism for miRNAs. [54] Ras-MEK regulation of the RNA-induced silencing complex (RISC) component Argonaute 2 (Ago2) has also been demonstrated to modify the miRNA content of sEVs. [55] However, there have been no previous reports demonstrating that H-Ras overexpression alone can significantly alter the proportions of small RNAs in EVs and preferentially load them with miRNAs. Considering that oncogenic activation of Ras isoforms is a tumor-initiating event in many types of cancers [56], the effect of H-Ras transformation on sEV RNA content was a significant observation in this study. Notably, the trend of increased miRNA content in sEVs continued with the progression of the MCF10A isogenic cell line series, with the greatest increase occurring at the transition from DCIS to IBC, indicating the involvement of sEV miRNA signaling in the progression of breast cancer from pre-invasive to invasive disease. Although this is a noteworthy in vitro observation, further investigations in additional in vivo models and in patients with breast cancer are warranted.

sEVs are known to signal through endocrine mechanisms by entering the vascular system and can therefore be detected in the blood plasma or serum. The content of circulating sEVs may serve as “tumor surrogates” or biomarkers for cancer progression and/or behavior. In this study, we used the MIND model to mimic the invasive progression of DCIS. sEV miRNAs that differed between DCIS and invasive cells, identified by RNA sequencing, were detected at varying levels in circulating plasma sEVs collected from mice with in situ and invasive cancers or with Rab27A knockdown. These data suggest that the transition from in situ to invasive cancer alters the release and/or levels of circulating sEV miRNAs, which reflects the behavior of the originating tumor cells. We previously reported that sEV miRNAs indicative of invasive cancer can be detected in plasma sEVs from patient-derived xenograft mice and human breast cancer patients. MiRNAs have been explored as biomarkers for predicting DCIS progression. [57] In our study, we observed increased levels of miR103a, miR-222, and miR-205 and reduced levels of miR-486 in the plasma of sEVs from mice with invasive cancers compared to those with DCIS. As no clinical biomarker currently exists to indicate DCIS to invasive progression, further follow-up studies using patient cohorts should be conducted to determine whether circulating sEVs miRNAs can serve as prognostic biomarkers of progression in patients diagnosed with DCIS.

sEVs play a well-established role in promoting cancer progression [58, 59]; however, the mechanisms by which sEV signaling contributes to the progression of pre-invasive breast cancer remain unclear. In this study, we investigated the role of sEV signaling in pre-invasive cancer progression in vivo and in vitro. In vivo, we demonstrated that the reduction in sEV secretion and depletion of sEV miRNAs diminished the number of invasive events. We specifically examined the effect of overexpressing miR-30, miR-103a, miR-205, and miR-222 on EMT, migration, and invasion and identified miR-205 as a potent inhibitor of invasion and EMT. Further examination of miR-205 in MCF10 progression series and invasive breast cancer cells demonstrated a consistent decrease in cellular levels of miR-205, with a concomitant increase in miR-205 in sEVs, which was corroborated by a significant reduction in miR-205 levels observed in primary breast tumors relative to normal tissues from large-scale patient datasets. Given the propensity for invasive cancer cells to preferentially release miR-205 into sEVs, we sought to examine how blocking the release of miR-205 via sEVs by Rab27A knockdown combined with over-expression of miR-205 would affect invasive programs in the cells. We demonstrated that either Rab27A knockdown or overexpression of miR-205 reduced proliferation and invasion and that combined Rab27A knockdown and miR-205 overexpression further reduced proliferation. Examination of gene expression programs altered by miR-205 overexpression in both wild-type and Rab27A knockdown cells revealed altered MAPK, EMT, ECM-receptor, and p53 signaling. Knockdown of Rab27A and miR-205 resulted in reduced TGFβ/SMAD signaling along with activation of stress-associated MAPK, p38, G1-cell cycle arrest, activation of p53, increased p21, early apoptosis, and necrosis. Collectively, these in vivo and in vitro experiments demonstrated that sEV miRNA signaling drives DCIS invasive progression and that inhibiting sEV miRNA release and increasing miR-205 levels results in reduced proliferation, EMT, and invasion, ultimately resulting in cell cycle arrest and cell death.

The mechanisms by which sEVs and specific sEV miRNAs promote breast cancer progression warrant further investigation. We speculate that the preferential release of miR-205 could function as a mechanism by which TGFβ becomes tumor promoting, by allowing TGFβ to proceed unregulated, resulting in a loss of sensitivity to the tumor suppressive role of TGFβ. Based on these observations, it is proposed that the combination of inhibiting sEV release and replenishing sEVs miRNAs could result in the cellular accumulation of miRNAs that can suppress EMT and pro-invasiveness, such as miR-205, leading to cellular stress and death. Further studies are necessary to examine this combinatorial effect using in vivo systems and the other sEV miRNAs identified in this study.

In conclusion, the present study demonstrates that the RNA content of sEVs undergoes progressive alterations during the stepwise transition from normal mammary epithelial cells to invasive breast cancer cells. The most significant changes in miRNA expression were observed during the transition from DCIS to invasive cancer. These alterations were also detected in circulating plasma sEVs in a mouse model of DCIS during invasive progression. Furthermore, sEVs and their miRNA content have been found to promote invasive programs through both autocrine and paracrine mechanisms. Inhibition of sEV release, combined with replenishment of their levels, results in stress responses, cell cycle arrest, and cell death.

## Ethics declarations

Animal experiments were conducted in accordance with guidelines and regulated under a protocol approved by the University of Kansas School of Medicine Animal Care and Use and Human Subjects Committee.

